# Fitness effects of CTX-M-15-encoding IncF plasmids on their native *Escherichia coli* ST131 *H30*Rx hosts

**DOI:** 10.1101/2021.08.17.456450

**Authors:** Jana Palkovicova, Iva Kutilova, Javier DelaFuente, Adam Valcek, Matej Medvecky, Ivana Jamborova, Ibrahim Bitar, Minh Duy Phan, Alvaro San Millan, Monika Dolejska

## Abstract

**Objectives:** The objective of this study was to investigate effects of large CTX-M-15-encoding IncF plasmids on the fitness of their native *E. coli* ST131 *H30*Rx hosts in order to understand possible plasmid-host coevolution.

**Methods:** We selected five *E. coli* ST131 *H30*Rx strains of diverse origin, each carrying a multireplicon IncF plasmid encoding the gene *bla*_CTX-M-15_. The plasmid was eliminated from each isolate by displacement using an incompatible plasmid vector pMDP5_cureEC958. Whole-genome sequencing (WGS) was performed to obtain complete chromosome and plasmid sequences of wild-type isolates and to detect chromosomal mutations in plasmid-free strains. Competition assays were conducted to determine the relative fitness of plasmid-free clones compared to the corresponding wild-type isolates.

**Results:** We were able to successfully eliminate the IncF plasmids from all of the wild-type strains using the curing vector pMDP5_cureEC958. The chromosomes of plasmid-free clones contained zero to six point mutations. Plasmid-free strains of three isolates showed no significant difference in relative fitness compared to the corresponding plasmid-free strains. In the two remaining isolates, the plasmids produced a small but significant fitness cost.

**Conclusion:** We conclude that IncF plasmids produce moderate fitness effects in their *E. coli* ST131 *H30*Rx hosts. This fitness compatibility is likely to promote the maintenance of antibiotic resistance in this worrisome *E. coli* lineage.

## Introduction

Extraintestinal pathogenic *E. coli* (ExPEC) represent a huge public health burden^1^ as these strains are a common source of numerous diseases, from mild to life-threatening infections, such as urinary tract or blood-stream associated infections, bacteraemia and meningitis^2^. Globally emerging multi-drug resistance, especially to antimicrobials of clinical importance, such as fluoroquinolones and cephalosporins, is of special concern in this species in recent years. Due to the resistance, only a few or even no therapeutic options are left for a treatment of infections caused by these bacteria^2,3^.

Resistance to clinically important antimicrobials among ExPEC strains was scarce before 2000. Since then, the number of ExPEC isolates resistant to fluoroquinolones and cephalosporins has been increasing exponentially^4^. Highly virulent and multi-drug resistant *E. coli* ST131 is one of the clinically most important ExPEC strains due to its worldwide predominance. Even though *E. coli* ST131 was mainly associated with human infections^5^, recent findings brought a disturbing evidence of its dissemination among companion animals, poultry, livestock, food products, wildlife and environment, including wastewater treatment plant effluents^6,7^.

A study from 2015^8^ analysed the evolution of *E. coli* ST131 lineage. Based on investigations from the mid-2000s, it is apparent that previous consumption and misuse of antimicrobials is linked with the emergence of resistant pathogens^8,9^. Antimicrobials, such as fluoroquinolones and cephalosporins, were often used for the treatment of human infections which resulted in the emergence of fluoroquinolone resistant subclone *E. coli* ST131 *H30*R. Furthermore, by acquisition of an incompatibility F (IncF) plasmid carrying an ESBL gene *bla*_CTX-M-15_, a distinct fluoroquinolone resistant extended-spectrum beta-lactamase (ESBL) producing *E. coli* ST131 subclone *H30*Rx emerged^9,10^. Compensatory mutations in combination with virulence determinants and resistance to clinically important antimicrobials allowed the subclone *H30*Rx to outcompete other sublineages of *E. coli* ST131 and in the late 2000s became a globally disseminated and most prevalent ExPEC subclone^8,11^.

IncF are complex epidemic resistance plasmids composed of more than one plasmid replicon. CTX-M-15-encoding IncF plasmids present in the subclone *H30*Rx usually harbour two plasmid replicons with plasmid multilocus sequence type (pMLST) F2:A1:B-. These plasmids typically harbour other resistance genes apart from *bla*_CTX-M-15_, such as *bla*_TEM-1_, *bla*_OXA-1_, *aac(6’)-Ib-cr, catB4, aadA5, mph*(A), *dfrA7, tet*(A) and *sul1*. Additionally, the narrow-host IncF plasmids encode partitioning and addiction systems to ensure their maintenance^6,8,12^. Carriage of such large plasmids providing selective advantage for a bacterial host via additional virulence and antibiotic determinants, usually imposes a fitness cost to its host^13^. On the other hand, a previous study suggested that *E. coli* ST131 *H30*Rx is adapted to large IncF plasmids^8^.

In this study, we analyse plasmid-host interactions between this intriguing *E. coli* subclone and its plasmids. We aimed to estimate the fitness impact of the large F2:A1:B-IncF plasmids, previously recognised as a foundation of *H30*Rx sublineage emergence, on its native host. Five representatives of the *E. coli* ST131 *H30*Rx were selected for elimination of IncF plasmid using the curing vector pMDP5_cureEC958. Plasmid fitness effects were subsequently calculated using competition assays between the plasmid-carrying and plasmid-free isogenic clones.

## Materials and Methods

### Bacterial strains

Five *E. coli* ST131 *H30*Rx strains were selected out of the collection of 169 *E. coli* ST131 isolates of diverse origin from several geographic regions^6^. Selected isolates were of human (n = 3), environmental (n = 1) and companion animal (n = 1) origin. Each strain carried a large IncF plasmid harbouring *bla*_CTX-M-15_. Additional information on isolates obtained during our previous study^6^ is presented in Table S1.

Bacterial strains were routinely grown on Luria-Bertani agar (LBA; Sigma-Aldrich, Saint Louis, USA) supplemented with cefotaxime (2 mg/L) at 37 °C overnight if not specified otherwise. Competition assays were performed in Luria-Bertani broth (LBB; Becton Dickinson, MD, USA).

### Construction of curing vector

The curing vector pMDP5_cureEC958 was designed based on the pCURE2 plasmid^14^. The backbone vector pMDP5 was assembled from three vectors including pUC19 (ori and MCS, 126-1480 nt), pKD3 (chloramphenicol resistant gene, 85-980 nt) and pCURE2 (*sacB* gene, 209-2076 nt). Subsequently, a curing fragment (containing RepFIIA, RepFIA, *ccdA, sok, pemI*, and *vagC*) was synthesised and cloned into pMDP5, creating the curing vector pMDP5_cureEC958 (Figure 1). All cloning and DNA synthesis steps were performed by Epoch Life Science (Texas, USA). The sequence of pMDP5_cureEC958 was deposited to GenBank under accession number MZ723317.

**Figure 1.**
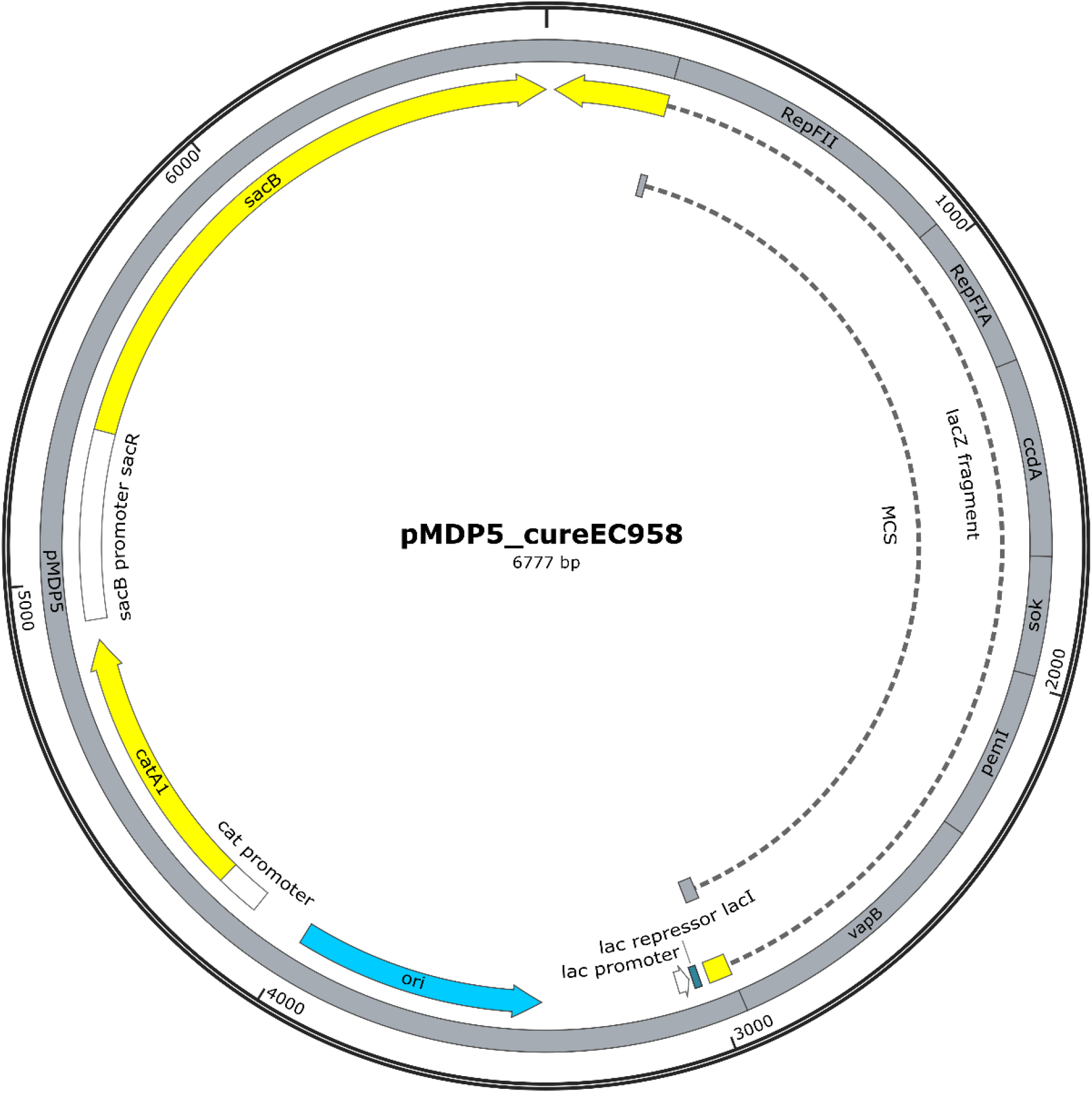
Genetic map of a plasmid vector pMDP5_cureEC958 designed for plasmid curing. It contains chloramphenicol resistance encoding gene *catA1* for selective cultivation of strains harbouring the vector and sucrose sensitivity gene *sacB* for selection of plasmid-free isolates disposed of the vector. For the purpose of plasmid curing, it harbours genes encoding antitoxins VapB, PemI, Sok and CcdA and IncF plasmid replicons RepFIA and RepFII.

### Plasmid curing

Plasmid-free variants were created using a curing method described by Hale and colleagues^14^ with the curing vector pMDP5_cureEC958. The plasmid curing vector was introduced into the wild-type strains by electroporation (1.8 kV, 25 µFar, 200 Ω) using Gene Pulser Xcell™ electroporation system (Bio-Rad Laboratories Inc., California, USA) as described before^15^. Cultures harbouring both, the wild-type plasmid and the designed construct, were selected on LBA supplemented chloramphenicol (30 mg/L) and the presence of *bla*_CTX-M_ and *catA1* genes was verified by PCR^16,17^. Transformants were cultivated on LBA containing chloramphenicol (30 mg/L) in order to eliminate the IncF plasmid. After successful removal of the wild-type plasmid, verified by the same PCR, the pMDP5_cureEC958 was eliminated on a non-selective LBA plates supplemented with 5% sucrose. Three to four plasmid-free clones of each isolate were selected.

Selected plasmid-free clones were sequenced on MiSeq platform (Illumina) as described below to investigate possible single nucleotide mutations (SNPs) on chromosomes. Subsequently, one plasmid-free clone of isolates without chromosomal mutations was selected to reintroduce the corresponding wild-type IncF plasmid as a control of experiment. Plasmid DNA was extracted from wild-type strains using Genopure Plasmid Midi Kit (Roche Diagnostics GmbH, Mannheim, Germany). Plasmids were reintroduced by electroporation and their presence was confirmed by PCR assays for gene *bla*_CTX-M_ and for FII and multiplex (FIA, FIB, FIC) IncF replicons^16,18^.

### Whole-genome sequencing

Wild-type isolates were subjected for short- and long-read sequencing. Additionally, plasmid-free strains and plasmid-free strains with reintroduced wild-type IncF plasmid were selected for short-read sequencing. Genomic DNA for short-read sequencing was extracted using NucleoSpin® Tissue kit (Macherey-Nagel, GmbH & Co. KG, Germany), library was prepared by Nextera® XT Library Preparation kit (Illumina, San Diego, CA, USA) and sequenced using 2×250 bp paired-end sequencing on MiSeq (Illumina) platform. NucleoSpin® Microbial DNA kit (Macherey-Nagel) was used for the extraction of genomic DNA aimed for long-read sequencing. Libraries were constructed using SMRTbell Express Template Prep Kit 2.0 (Pacific Biosciences, PacBio, USA) followed by single molecule real-time (SMRT) sequencing on Sequel I Platform (PacBio).

### Data analysis

Raw reads acquired by Illumina sequencing were trimmed using Trimmomatic v0.39^19^ to remove adaptor residues and discard low quality read regions (Q ≤ 20). SPAdes v3.13.1^20^ with the “--careful” configuration was used to obtain *de novo* assemblies. Center for Genomic Epidemiology tools (PlasmidFinder v2.1, pMLST v2.0, ResFinder v4.0) were used to verify the presence of plasmid replicons and genes intermediating antibiotic resistance (https://cge.cbs.dtu.dk/services/). HGAP4 in SMRT Link v.6 (PacBio) was used to obtain polished long reads in fastq format. Hybrid assembly of trimmed short and long reads was performed using Unicycler v0.4.8^21^ and corrected with Pilon v1.23^22^ in order to reconstruct chromosome and plasmid sequences of wild-type isolates. Complete circular sequences of plasmids were manually annotated using Geneious v7.1.9 (Biomatters, Auckland, New Zealand) in compliance with annotation form of previous studies^23^.

### Comparative genomics

Phylogenetic relatedness of wild-type isolates was estimated. Prokka v1.14.1^24^ was used to predict open reading frames of isolates assemblies and their core genome was aligned using Roary v3.12.0^25^. Subsequently, the alignment was used to generate phylogenetic tree in RAxML v8.2.11^26^ under GTR+GAMMA model supported by 1,000 bootstraps. A nucleotide similarity between the isolates was estimated using the core genome alignment in snp-dists v0.6.3 (https://github.com/tseemann/snp-dists) considering the number of SNPs. The phylogenetic tree was visualized in iTOL v5.7^27^.

BLAST (Basic Local Alignment Search Tool) of NCBI (National Center for Biotechnology Information, MD, USA) was used to find and download a plasmid sequence with the highest coverage and identity from GenBank. Genetic content of the IncF plasmids was compared using BLAST Ring Image Generator (BRIG) v0.95^28^ and Clinker v0.0.13^29^. Presence and nomenclature of specific insertion sequences in IncF plasmids were confirmed using ISfinder database^30^ and toxin-antitoxin systems were verified using Conserved Domain Database^31^.

Comparison of the wild-type isolates to the corresponding plasmid-free strains and to the plasmid-free strains with reintroduced plasmids was made to verify the identity of the strains as well as the identity of the wild-type and reintroduced plasmids. Corresponding sequences were aligned using algorithm BWA-MEM v0.7.17^32^ and manually checked in Geneious v7.1.9.

### Single nucleotide polymorphism analysis

Corrected short reads of plasmid-free variants were mapped to the corresponding wild-type *de novo* assemblies using Bowtie2 v2.3.5^33^. Variant calling was performed by VarScan v2.4.4^34^ based on the coverage of mapped reads. Minimum variant frequency was 80% and called variants were manually checked in Geneious v7.1.9. Corresponding wild-type reads were mapped and analysed as well in order to normalize obtained results.

### Transferability of IncF plasmids

Wild-type *E. coli* ST131 *H30*Rx isolates were used as donors while laboratory strain *E. coli* TOP10 (Invitrogen Life Technologies, Carlsbad, CA, USA) and corresponding plasmid-free variants of studied isolates were used as recipients for the estimation of conjugation ability of IncF plasmids using filter mating assays based on a previous study^35^. Conjugations were conducted in technical triplicates and biological duplicates.

### Relative fitness measurements

Competition assays were performed to compare the relative fitness of the wild-type strains and their plasmid-free clones using flow cytometry as previously described^36^. Only the plasmid-free strains without chromosomal mutations were selected for fitness experiments. A small non-mobilisable pBGC plasmid (MT702881)^37^ producing green fluorescent protein (GFP) was transformed to the wild-type strains by electroporation^15^. Transformants were selected on LBA plates containing cefotaxime (2 mg/L) and chloramphenicol (30 mg/L) and subsequently verified by PCR assays for genes *bla*_CTX-M_ and *gfp*^16,37^.

Two competition assays, each consisting of six biological replicates, were performed for each isolate. Competitions were performed between GFP-tagged wild-type strains and their untagged plasmid-free variants while each included a competition between tagged and untagged wild-types for normalisation. Overnight cultures were mixed in ratio 1:1 and diluted 1:400 for the competition. GFP expression in the wild-type strains resulting in fluorescence was induced by incubation in 0.9% sodium chloride solution containing 0.5% L-arabinose for 1.5 hours. Plasmid-free and wild-type populations were competed at 37 °C for 22 hours shaking at 225 rpm. Initial and final proportions were measured on NovoCyte (ACEA Biosciences) flow cytometer recording 50,000 events of each mixture. Relative fitness of plasmid-free clones compared to corresponding wild-types was estimated using the formula:

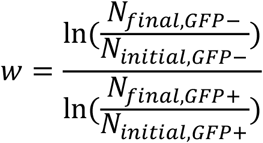

where *w* represents relative fitness, N_initial, GFP-_ and N_final, GFP-_ are initial and final values of untagged population and N_initial, GFP+_ and N_final, GFP+_ are proportions of GFP-marked population before and after competition. Relative fitness of plasmid-free clones was statistically processed using Student’s T-test where relative fitness with *p* value < 0.05 was evaluated as statistically significant. Obtained data was normalized using a competition between the tagged and untagged wild-type populations in order to capture relative fitness of plasmid-free clones in comparison to the corresponding (untagged) wild-type isolates. Competitions between wild-type strains and constructed plasmid-free strains with reintroduced wild-type IncF plasmid were performed as a control.

## Results

### Strain and plasmid features

In order to compare fitness effects of the plasmid on its native host, *E. coli* ST131 *H30*Rx isolates carrying a single large *bla*_CTX-M-15_ harbouring IncF plasmid were selected. The phylogenetic analysis of the five selected strains (Table S1) was based on the core-genome alignment of 4,803 genes and showed 78-440 SNPs differences (Figure S1).

All five plasmids contained two IncF replicons (RepFIA, RepFII) with pMLST formula F2:A1:B-, slightly varied in size and antibiotic resistance genes content (Figure S2). All plasmids provided multi-drug resistance profile, encoding genes for ESBL as well as for other antimicrobials and contained insertion sequences, mostly IS*26* (Table 1). The *ccdAB* and *pemIK* toxin-antitoxin systems were encoded in all plasmids within replicons RepFIA and RepFII, respectively. Each plasmid harboured two copies of the addiction system *vapBC*. All but one plasmid (pM45) harboured *hok/sok* system and plasmids of human isolates encoded *parDE* system.

**Table 1.**
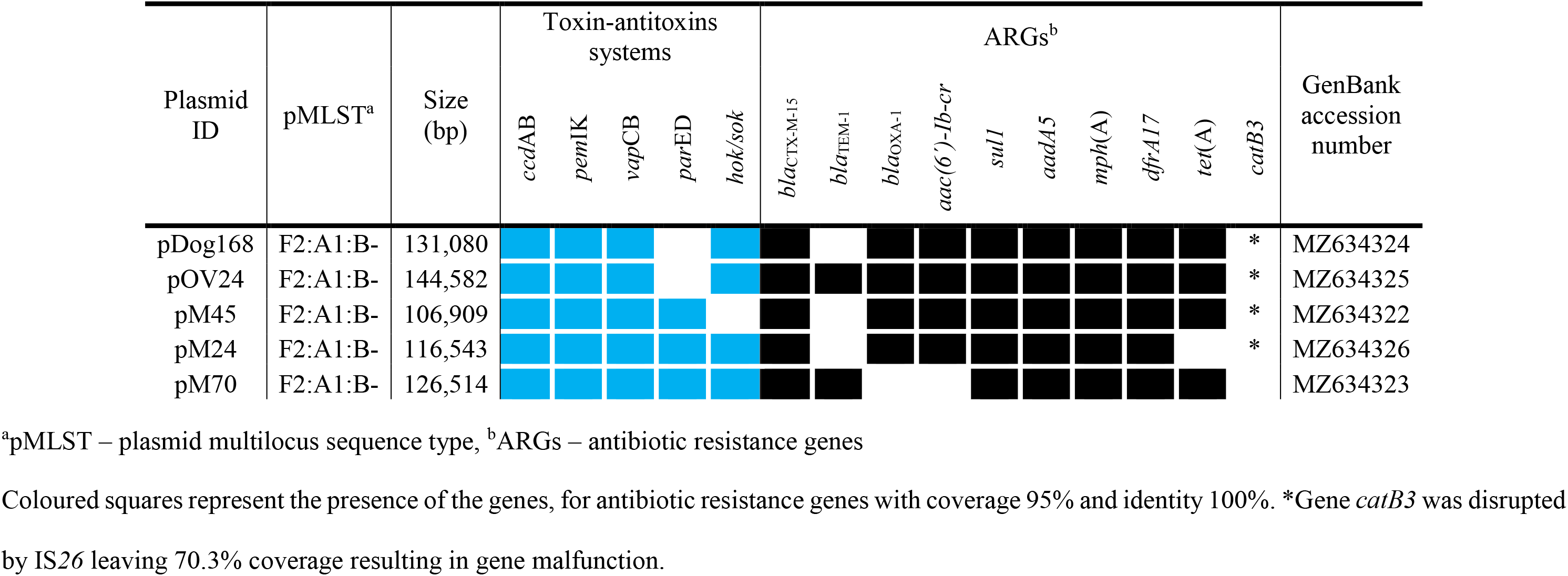
Selected genetic characteristics of CTX-M-15-encoding IncF plasmids in our study.

All IncF plasmids in our study have proved to be non-conjugative. Genetic analysis of transfer (*tra*) regions showed diverse defects in all plasmids likely resulting in their non-functionality (Figure 2). The *tra* region of pOV24 was disrupted in two parts by the *bla*_TEM-1_ gene, usually transposed within a composite mobile genetic element, but 3’ flanking sequence IS*26* was disrupted by IS*Ecp1* element. Furthermore, the *traC* gene was truncated by another IS*26* and genes *traW* and *traU*, encoding proteins for pilus assembly and DNA transfer, were missing. The *tra* region of pM70 was disrupted by composite mobile element containing *bla*_TEM-1_ gene flanked by IS*15DI* and IS*26* similarly as in pOV24. Moreover, part of the second half of the *tra* genes was translocated 29.5 kb from the first part of the *tra* region and truncated by IS*26* in *traN* gene. The first part of the *tra* region, including *traJ*, serving as a transcriptional activator of the *tra* region, was completely missing in plasmids pM24 and pM45. Plasmid pDog168 lacked most of this part of the region as well with an exception of the *traM* gene.

**Figure 2.**
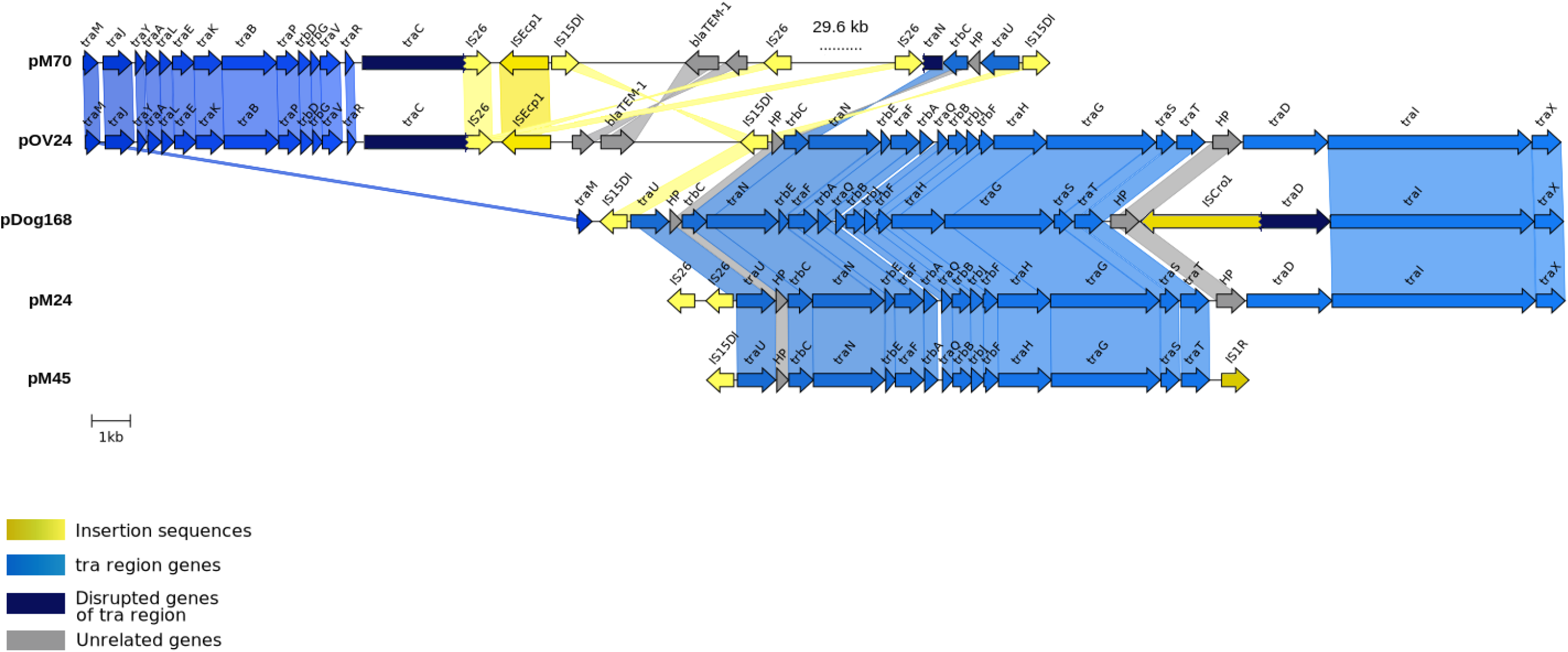
Transfer region of studied plasmids. The shading shows the similarity above 99.9%.

### Plasmid curing

To study plasmid-associated fitness effects on their native host, all five wild-type strains were cured of the naturally occurring IncF plasmids. A curing vector pMDP5_cureEC958 was designed for the generation of plasmid-free variants, harbouring selective genes (*catA1* and *sacB*), replicons RepFIA, RepFII, and antitoxins of the addiction systems encoded by wild-type plasmids (Figure 1).

Four plasmid-free clones per each of four isolates (M24, M45, M70 and Dog168) and all three grown plasmid-free clones of the isolate OV24 were selected for further analyses. Sequence comparison of plasmid-free clones to the corresponding wild-type isolates discovered zero to six chromosomal mutations. Mutations occurred in 52.6% of plasmid-free strains (in 10 out of 19). Nearly all mutations (92%, 23/25) occurred in protein coding sequences, only two of them were located in intergenic regions. Additionally, most of the mutations in coding sequences (82.6%, 19/23) were non-synonymous. No mutations occurred only in one of the plasmid-free clones of the OV24 (1/3) and M70 (1/4) isolates, in two Dog168 (2/4), two M24 (2/4) and three M45 (3/4) plasmid-free clones. Detailed list of genetic changes in plasmid-free clones is in Table S2.

Four of all plasmid-free strains with the reintroduced wild-type plasmid (40%, 4/10) harboured one mutation. Although, only one of them was non-synonymous.

### Fitness impact of IncF plasmids on their native host

To maintain isogenic conditions in the competition experiment, only the plasmid-free clones with no mutations in their chromosome were selected for the measurement of the IncF-associated fitness effects on their native host. Relative fitness of plasmid-free clones was estimated in comparison to the corresponding wild-type isolates considering a background fitness of wild-types as 1 (Table 2). Analysis of two competition assays both consisting of six biological replicates for each combination plasmid-free clone and wild-type revealed non-significant fitness effects (*p* > 0.05) of IncF plasmids in three isolates (Dog168, OV24, M45). Moderate increase (*p* < 0.05) of relative fitness was observed in plasmid-free strains of two isolates (M24 and M70) as visualised in Figure 3, revealing a small plasmid fitness cost.

**Table 2.** Relative fitness of the plasmid-free clones in comparison to their wild-type isolates

| Isolate ID | Plasmid-free clone ID<sup>a</sup> | Relative fitness<sup>b</sup> (±SD) | *p* value |
| --- | --- | --- | --- |
| Dog168 | CC4 | 0.992 ± 0.06 | 0.764 |
| | CC5 | 1.025 ± 0.03 | 0.216 |
| OV24 | CC2 | 1.015 ± 0.04 | 0.284 |
| M45 | CC2 | 0.985 ± 0.04 | 0.386 |
| | CC3 | 1.012 ± 0.03 | 0.444 |
| | CC4 | 1.024 ± 0.03 | 0.108 |
| **M24** | **CC5** | **1.038 ± 0.02** | **1.8 x 10<sup>-6</sup>** |
|  | **CC8** | **1.021 ± 0.01** | **4 x 10<sup>-4</sup>** |
| **M70** | **CC1** | **1.027 ± 0.02** | **1.5 x 10<sup>-3</sup>** |

<sup>a</sup> ID of constructed plasmid-free strains; CC stands for cured clone, <sup>b</sup> Relative fitness of plasmid- free strain compared to the corresponding wild-type isolate which background fitness was estimated as 1. Isolates highlighted in bold showed significant ( $p < 0.05$ ) relative fitness changes, however, the increase was moderate.

**Figure 3.**
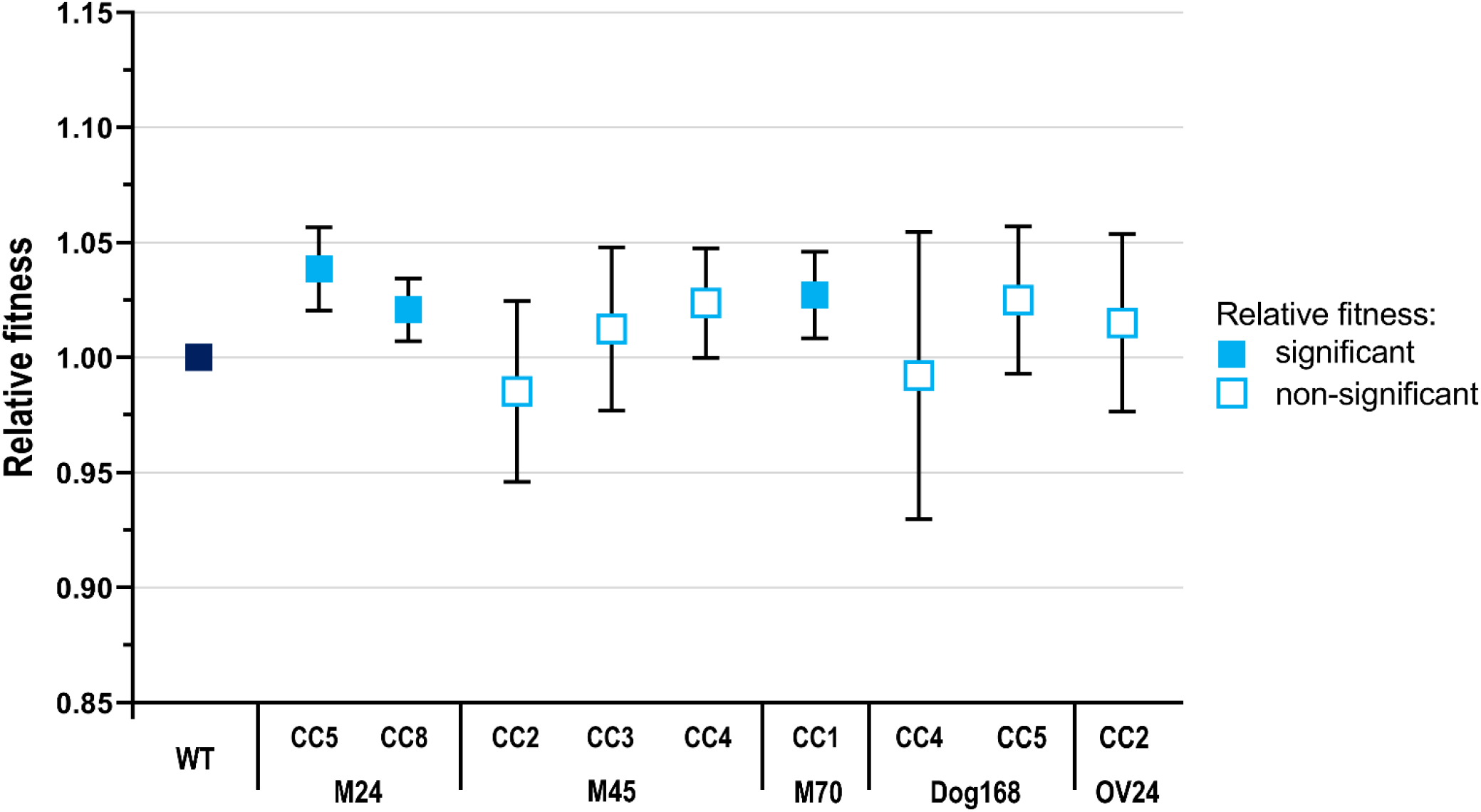
Relative fitness of plasmid-free clones in comparison to the corresponding wild-type isolates. Background fitness of wild-types was estimated as 1. CC stands for cured clone. Bars indicate standard deviation.

Relative fitness differences of the two M24 plasmid-free clones were statistically significant (with *p*-value 1.83 × 10^−6^ and 4.07 × 10^−4^, respectively), however, the increase in relative fitness was moderate (1.04 and 1.02). A similar result was observed for the M70 plasmid-free clone, which showed a moderate but significant increase in relative fitness compared to the wild type clone (*w* = 1.03, *p* = 1.5 × 10^−3^).

## Discussion

Even though the *E. coli* ST131 has attracted much attention in the last few years due to its predominance in ExPEC infections, its success is still not fully elucidated. Plasmids play a key role in bacterial survival under a selection pressure by providing virulence and antibiotic resistance genes, but it is known that they often impose a fitness burden to their hosts^38–40^. However, the fitness cost of a specific plasmid can differ in various hosts^41^.

In our study, we investigated the fitness effects of strictly clade-specific^12^ F2:A1:B-IncF plasmids, previously recognised as a source of *H30*Rx subclone emergence^8,11^, on these native hosts. Five native *E. coli* ST131 *H30*Rx hosts were cured of the large *bla*_CTX-M-15_-harbouring IncF plasmids and then competed against their corresponding wild-type strain to calculate the plasmid fitness impact.

### Plasmid curing

Plasmid curing was previously recognised as a best way to study effects of a plasmid on a bacterial population^38^. Traditionally used methods for plasmid curing involved bacterial growth in the presence of a chemical factor^42^. Although these techniques were widely used, efficiency of plasmid curing was low and promoted high accumulation of unwanted mutations^43^. Recently, the CRISPR (Clustered Regularly Interspaced Short Palindromic Repeats) curing is being used more often. The CRISPR curing is based on targeting a specific conserved sequence within plasmids and subsequent plasmid cleavage. This technique is efficient and do not generate new mutations, however, there are still plasmids which avoid targeting^44^. As IncF plasmids are known for their complexity and many combinations of the replicon alleles among the plasmids, the pCURE method^14^ with the vector pMDP5_cureEC958 was used in this study. The method is based on incompatibility of the targeted plasmid with an introduced curing vector. Even though many plasmids overcome incompatibility elimination, it is possible to design a construct with more replicons and antitoxin genes to successfully cure a strain of a plasmid. The approach proved efficient in the past^14,44^ as well as in our study. On the other hand, we observed point mutation accumulation in 52.6% (10 out of 19) of the plasmid-free clones. Therefore, WGS after pCURE plasmid curing followed by genomic comparison of plasmid-free clones and their corresponding wild-type isolates is necessary to exclude the clones harbouring mutations and to obtain reliable relative fitness results.

### Fitness effects of the F2:A1:B-plasmids

The fitness cost imposed by plasmids is influenced by several factors. It was observed before, that solely the size of plasmids plays no role in their fitness cost, however, maintaining the plasmid-encoded accessory genes can produce an energetic burden. The increasing number of accessory genes, such as antimicrobial resistance genes, correlates with the higher fitness cost^45^.

In order to evaluate the fitness impact of the F2:A1:B-plasmids providing their hosts with multi-drug resistance, we estimated relative fitness of the plasmid-free strains in comparison to the corresponding wild-type isolates. Competitions followed by measurement using flow cytometer were used for this purpose. This method is considered much more sensitive than growth curve analysis allowing to detect even subtle differences in relative fitness^13^. We demonstrated that fitness impact of these IncF plasmids on their native hosts in non-selective conditions was moderate to negligible in human as well as in animal and environmental isolates which is in concordance with previous studies on F2:A1:B-plasmids^46–48^. A study of Shin and Ko^47^ focused on effects of CTX-M-14 and CTX-M-15 IncF plasmids from human clinical isolates and their impact on a laboratory *E. coli* J53 strain. Based on the growth curve analysis of the transconjugants, the authors proposed that strains harbouring *bla*_CTX-M_ on IncF plasmids were as competitive as the naive host. Ranjan and his colleagues^48^ studied competitiveness of *E. coli* ST131 harbouring IncF plasmids and their plasmid-free variants against colicin-producing *E. coli* ST10. The authors observed similar fitness (*p* > 0.05) of the wild-type *E. coli* ST131 and their plasmid-free variants based on growth curves analysis. However, growth curves in this experiment were conducted on selective plates which could affect the fitness of the strains as some antibiotic resistance genes are genetically linked to each other and co-selected. Therefore, the supplementation of plates with antibiotics could create a selection pressure where carriage of plasmids would be more beneficial for the strain survival^49^. Mahérault and colleagues^46^ provided the investigation on two human clinical *bla*_CTX-M-15_-harbouring F2:A1:B-plasmids and their impact on an *E. coli* J53-2. Even though the fitness cost of an IncF plasmid occurred initially after conjugation, the authors observed that this fitness cost alleviated and a transconjugant carrying the IncF plasmid proved more competitive than a transconjugant harbouring an IncC plasmid.

Previous studies also indicated that a functional conjugation system could have a negative impact on a bacterial fitness and plasmids use several different ways to supress the conjugative transfer^39^. Even though the silencing of the conjugative transfer results in a decrease of horizontal spread of the plasmids, the vertical spread is supported by a fitness cost reduction^50^. The *tra* region responsible for conjugative transfer investigated during our study was incomplete, the missing genes and length of the missing sequences differed among studied plasmids. The rearrangements resulted in the non-functionality of the *tra* region of each plasmid. Conjugation malfunction together with plasmid addiction systems ensure the vertical transmission of the IncF plasmids. Additionally, it was pointed out that the initial fitness cost is reduced over time. This phenomenon was observed in long-term evolution experiments, even though the fitness of the plasmid-bearing strains was lower than of those without plasmids in many cases^39^.

We demonstrated that fitness impact of these IncF plasmids on their native hosts in non-selective conditions was moderate to negligible among phylogenetically unrelated isolates of diverse origin. Our results, combined with previous findings, strongly suggest that *E. coli H30*Rx and IncF plasmids form successful associations promoting the world-wide dissemination of this ExPEC lineage.

## Acknowledgements

We thank Jarka Lausova and Dana Cervinkova from University of Veterinary Sciences Brno (Czech Republic) and Aida Alonso del Valle and Carmen de la Vega from Hospital Universitario Ramon y Cajal (Spain) for their work in the laboratory. Furthermore, we are grateful to Kristina Nesporova from University of Veterinary Sciences Brno and Jaroslav Hrabak from the University Hospital Pilsen (Czech Republic) for the help with short- and long-read sequencing, respectively. We also thank Chris Thomas for the inspiration and help in designing the curing plasmid.

## Transparency declarations

None to declare.

## References

1. World Health Organization. Antimicrobial Resistance: Global Report on Surveillance. Geneva, Switzerland; 2014.

2. Schwaber MJ, Navon-Venezia S, Kaye KS, Ben-Ami R, Schwartz D, Carmeli Y. Clinical and Economic Impact of Bacteremia with Extended-Spectrum-β-Lactamase-Producing Enterobacteriaceae. Antimicrob Agents Chemother 2006; 50: 1257–62.

3. Mathers AJ, Peirano G, Pitout JDD. Escherichia coli ST131: The Quintessential Example of an International Multiresistant High-Risk Clone. In: Advances in Applied Microbiology. Vol 90. Elsevier Ltd, 2015; 109–54.

4. Pitout JDD. Extraintestinal pathogenic Escherichia coli: an update on antimicrobial resistance, laboratory diagnosis and treatment. Expert Rev Anti Infect Ther 2012; 10: 1165–76.

5. Poolman JT, Wacker M. Extraintestinal Pathogenic Escherichia coli, a Common Human Pathogen: Challenges for Vaccine Development and Progress in the Field. J Infect Dis 2016; 213: 6–13.

6. Jamborova I, Johnston BD, Papousek I, et al. Extensive Genetic Commonality among Wildlife, Wastewater, Community, and Nosocomial Isolates of Escherichia coli Sequence Type 131 (H30R1 and H30Rx Subclones) That Carry bla_CTX-M-27_ or bla_CTX-M-15_. Antimicrob Agents Chemother 2018; 62: e00519–18.

7. Kidsley AK, White RT, Beatson SA, et al. Companion Animals Are Spillover Hosts of the Multidrug-Resistant Human Extraintestinal Escherichia coli Pandemic Clones ST131 and ST1193. Front Microbiol 2020; 11: 1968.

8. Mathers AJ, Peirano G, Pitout JDD. The role of epidemic resistance plasmids and international high-risk clones in the spread of multidrug-resistant Enterobacteriaceae. Clin Microbiol Rev 2015; 28: 565–91.

9. Ben Zakour NL, Alsheikh-Hussain AS, Ashcroft MM, et al. Sequential acquisition of virulence and fluoroquinolone resistance has shaped the evolution of Escherichia coli ST131. mBio 2016; 7: e00347–16.

10. Riley LW. Pandemic lineages of extraintestinal pathogenic Escherichia coli. Clin Microbiol Infect 2014; 20: 380–90.

11. Price LB, Johnson JR, Aziz M, et al. The epidemic of Extended-Spectrum-β-Lactamase-Producing Escherichia coli ST131 Is Driven by a Single Highly Pathogenic Subclone, H30-Rx. mBio 2013; 4: e00377–13.

12. Kondratyeva K, Salmon-Divon M, Navon-Venezia S. Meta-analysis of Pandemic Escherichia coli ST131 Plasmidome Proves Restricted Plasmid-clade Associations. Sci Rep 2020; 10: 36.

13. San Millan A. Evolution of Plasmid-Mediated Antibiotic Resistance in the Clinical Context. Trends Microbiol 2018; 26: 978–85.

14. Hale L, Lazos O, Haines AS, Thomas CM. An efficient stress-free strategy to displace stable bacterial plasmids. Biotechniques 2010; 48: 223–8.

15. Wu N, Matand K, Kebede B, Acquaah G, Williams S. Enhancing DNA electrotransformation efficiency in Escherichia coli DH10B electrocompetent cells. Electron J Biotechnol 2010; 13.

16. Lewis II JS, Herrera M, Wickes B, Patterson JE, Jorgensen JH. First report of the Emergence of CTX-M-type Extended-Spectrum β-Lactamases (ESBLs) as the Predominant ESBL Isolated in a U.S. Health Care System. Antimicrob Agents Chemother 2007; 51: 4015–21.

17. Faldynova M, Pravcova M, Sisak F, et al. Evolution of Antibiotic Resistance in Salmonella enterica Serovar Typhimurium Strains Isolated in the Czech Republic between 1984 and 2002. Antimicrob Agents Chemother 2003; 47: 2002–5.

18. Carattoli A. Resistance plasmid families in Enterobacteriaceae. Antimicrob Agents Chemother 2009; 53: 2227–38.

19. Bolger AM, Lohse M, Usadel B. Trimmomatic: A flexible trimmer for Illumina sequence data. Bioinformatics 2014; 30: 2114–20.

20. Bankevich A, Nurk S, Antipov D, et al. SPAdes: A new genome assembly algorithm and its applications to single-cell sequencing. J Comput Biol 2012; 19: 455–77.

21. Wick RR, Judd LM, Gorrie CL, Holt KE. Unicycler: Resolving bacterial genome assemblies from short and long sequencing reads. PLoS Comput Biol 2017; 13: e1005595.

22. Walker BJ, Abeel T, Shea T, et al. Pilon: An integrated Tool for Comprehensive Microbial Variant Detection and Genome Assembly Improvement. PLoS One 2014; 9: e112963.

23. Papagiannitsis CC, Kutilova I, Medvecky M, Hrabak J, Dolejska M. Characterization of the Complete Nucleotide Sequences of IncA/C2 Plasmids Carrying In809-Like Integrons from Enterobacteriaceae Isolates of Wildlife Origin. Antimicrob Agents Chemother 2017; 61: e01093–17.

24. Seemann T. Prokka: rapid prokaryotic genome annotation. Bioinformatics 2014; 30: 2068–9.

25. Page AJ, Cummins CA, Hunt M, et al. Roary: rapid large-scale prokaryote pan genome analysis. Bioinformatics 2015; 31: 3691–3.

26. Stamatakis A. RAxML version 8: a tool for phylogenetic analysis and post-analysis of large phylogenies. Bioinformatics 2014; 30: 1312–3.

27. Letunic I, Bork P. Interactive Tree of Life (iTOL) v4: Recent updates and new developments. Nucleic Acids Res 2019; 47: 256–9.

28. Alikhan NF, Petty NK, Ben Zakour NL, Beatson SA. BLAST Ring Image Generator (BRIG): Simple prokaryote genome comparisons. BMC Genomics 2011; 12: 402.

29. Gilchrist CLM, Chooi Y-H. clinker & clustermap.js: automatic generation of gene cluster comparison figures. Bioinformatics 2021: btab007.

30. Siguier P, Perochon J, Lestrade L, Mahillon J, Chandler M. ISfinder: the reference centre for bacterial insertion sequences. Nucleic Acids Res 2006; 34: D32–6.

31. Marchler-Bauer A, Derbyshire MK, Gonzales NR, et al. CDD: NCBI’s conserved domain database. Nucleic Acids Res 2015; 43: D222–6.

32. Li H. Aligning sequence reads, clone sequences and assembly contigs with BWA-MEM. 2013: 1–3.

33. Langmead B, Trapnell C, Pop M, Salzberg SL. Ultrafast and memory-efficient alignment of short DNA sequences to the human genome. Genome Biol 2009; 10: R25.

34. Koboldt DC, Chen K, Wylie T, et al. VarScan: variant detection in massively parallel sequencing of individual and pooled samples. Bioinformatics 2009; 25: 2283–5.

35. Top E, Vanrolleghem P, Mergeay M, Verstraete W. Determination of the Mechanism of Retrotransfer by Mechanistic Mathematical Modeling. J Bacteriol 1992; 174: 5953–60.

36. DelaFuente J, Rodriguez-Beltran J, San Millan A. Methods to Study Fitness and Compensatory Adaptation in Plasmid-Carrying Bacteria. In: Horizontal Gene Transfer. Methods in Molecular Biology. Vol 2075. New York: Humana, 2020; 371–82.

37. Alonso-del Valle A, León-Sampedro R, Rodríguez-Beltrán J, et al. Variability of plasmid fitness effects contributes to plasmid persistence in bacterial communities. Nat Commun 2021; 12.

38. San Millan A, Maclean RC. Fitness Costs of Plasmids: a Limit to Plasmid Transmission. Microbiol Spectr 2017; 5: MTBP-0016-2017.

39. Porse A, Schønning K, Munck C, Sommer MOA. Survival and Evolution of a Large Multidrug Resistance Plasmid in New Clinical Bacterial Hosts. Mol Biol Evol 2016; 33: 2860–73.

40. San Millan A, Peña-Miller R, Toll-Riera M, et al. Positive selection and compensatory adaptation interact to stabilize non-transmissible plasmids. Nat Commun 2014; 5: 5208.

41. Johnson TJ, Singer RS, Isaacson RE, et al. In Vivo Transmission of an IncA/C Plasmid in Escherichia coli Depends on Tetracycline Concentration, and Acquisition of the Plasmid Results in a Variable Cost of Fitness. Appl Environ Microbiol 2015; 81: 3561–70.

42. Zaman M, Pasha M, Akhter M. Plasmid Curing of Escherichia coli Cells with Ethidium Bromide, Sodium Dodecyl Sulfate and Acridine Orange. Bangladesh J Microbiol 2010; 27: 28–31.

43. Lauritsen I, Porse A, Sommer MOA, Nørholm MHH. A versatile one-step CRISPR-Cas9 based approach to plasmid-curing. Microb Cell Fact 2017; 16: 135.

44. Li Y, Lin Z, Huang C, et al. Metabolic engineering of Escherichia coli using CRISPR-Cas9 meditated genome editing. Metab Eng 2015; 31: 13–21.

45. Vogwill T, Maclean RC. The genetic basis of the fitness costs of antimicrobial resistance: a meta-analysis approach. Evol Appl 2015; 8: 284–95.

46. Mahérault A-C, Kemble H, Magnan M, et al. Advantage of the F2:A1:B-IncF Pandemic Plasmid over IncC Plasmids in In Vitro Acquisition and Evolution of bla_CTX-M_ Gene-Bearing Plasmids in Escherichia coli. Antimicrob Agents Chemother 2019; 63: e01130–19.

47. Shin J, Ko KS. Effect of plasmids harbouring bla_CTX-M_ on the virulence and fitness of Escherichia coli ST131 isolates. Int J Antimicrob Agents 2015; 46: 214–8.

48. Ranjan A, Scholz J, Semmler T, et al. ESBL-plasmid carriage in E. coli enhances in vitro bacterial competition fitness and serum resistance in some strains of pandemic sequence types without overall fitness cost. Gut Pathog 2018; 10: 24.

49. Andersson DI, Hughes D. Antibiotic resistance and its cost: Is it possible to reverse resistance? Nat Rev Microbiol 2010; 8: 260–71.

50. Fernandez-Lopez R, del Campo I, Revilla C, Cuevas A, de la Cruz F. Negative Feedback and Transcriptional Overshooting in a Regulatory Network for Horizontal Gene Transfer. PLoS Genet 2014; 10: e1004171.

