## Supplementary Materials for "Fitness effects of CTX-M-15-encoding IncF plasmids on their native *Escherichia coli* ST131 *H30*Rx hosts"

### SUPPLEMENTARY MATERIAL

**Figure S1** Phylogenetic tree of the *E. coli* ST131 *H30*Rx representatives selected for this study.

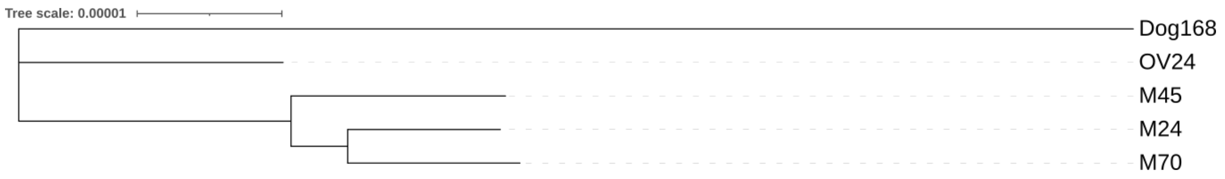

**Figure S2** Genetic comparison of IncF plasmids from this study with a similar plasmid (CP013657) retrieved from GenBank.

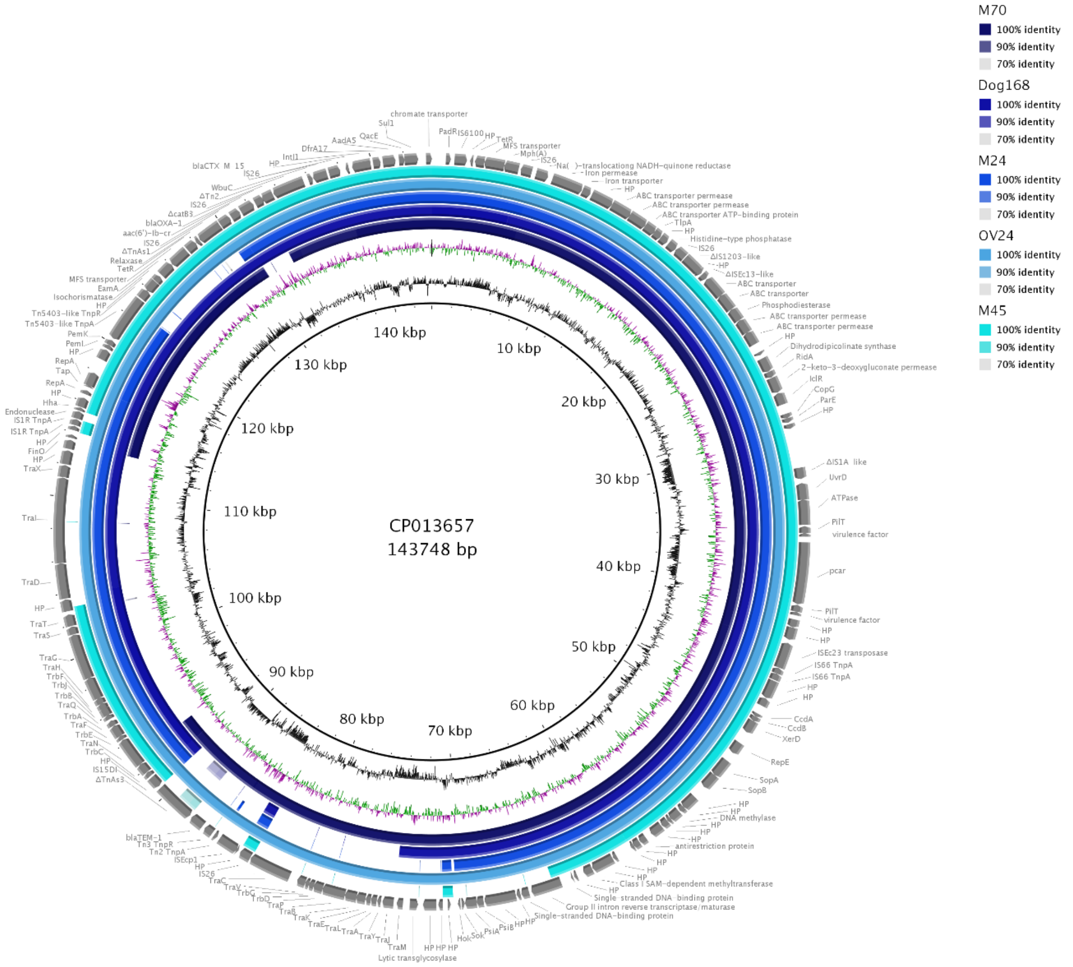

**Table S1** Characteristics of selected *E. coli* ST131 *H30*Rx isolates obtained during our previous study[1]

| Isolate ID | Sampling year | Origin[a] | | | PFGE | | Antibiotic resistance profile[b] | | | | | | | | | | | | | | | | | | | | | | | |
|---|---|---|---|---|---|---|---|---|---|---|---|---|---|---|---|---|---|---|---|---|---|---|---|---|---|---|---|---|---|---|
| | | Human | WWTP | dog | cluster | type | AMP | PIP | CFZ | CXM | CAZ | CTX | CFP | FEP | ATM | AMC | SAM | TZP | NAL | OFX | CIP | SXT | TET | DOX | SUL | TMP | AZM | STR | TOB | AMK |
| Dog168 | 2009 | | | ■ | 16 | 812 | ■ | ■ | ■ | ■ | | ■ | | | | ■ | ■ | | ■ | ■ | ■ | ■ | ■ | ■ | ■ | ■ | ■ | ■ | ■ | |
| OV24 | 2009 | | ■ | | 16 | 812 | ■ | ■ | ■ | ■ | | ■ | | | ■ | ■ | | | ■ | ■ | ■ | ■ | ■ | ■ | ■ | ■ | ■ | ■ | ■ | |
| M45 | 2010 | ■ | | | 16 | 1676 | ■ | ■ | ■ | ■ | ■ | ■ | ■ | ■ | ■ | ■ | ■ | | ■ | ■ | ■ | ■ | ■ | ■ | ■ | ■ | ■ | ■ | ■ | |
| M24 | 2009 | ■ | | | 23 | 1735 | ■ | ■ | ■ | ■ | ■ | ■ | ■ | ■ | ■ | ■ | ■ | ■ | ■ | ■ | ■ | ■ | ■ | | ■ | ■ | ■ | ■ | ■ | ■ |
| M70 | 2010 | ■ | | | 23 | 1735 | ■ | ■ | ■ | ■ | | ■ | ■ | | ■ | ■ | ■ | | ■ | ■ | ■ | ■ | ■ | ■ | ■ | ■ | ■ | ■ | | |

[a]Black squares indicate the origin of the isolate. Human isolates originated from urinary tract infections, St. Anne's Faculty Hospital in Brno, Czech Republic; WWTP – effluent of wastewater treatment plant Modrice Brno, Czech Republic; dog sample originated in Kenya

[b]Black squares indicate the resistance to the antibiotics. None of the isolates was resistant to meropenem, ertapenem, cefoxitin, gentamicin, nitrofurantoin, tigecycline, chloramphenicol and colistin; Ampicillin (AMP), Piperacillin (PIP), Cefazolin (CFZ), Cefuroxime (CXM), Ceftazidime (CAZ), Cefotaxime (CTX), Cefoperazone (CFP), Cefepime (FEP), Aztreonam (ATM), Amoxicillin-clavulanic acid (AMC), Amplicillin/sulbactam (SAM), Piperacillin/tazobactam (TZP), Nalidixic acid (NAL), Ofloxacin (OFX), Ciprofloxacin (CIP), Trimethoprim/sulfamethoxazole (SXT), Tetracycline (TET), Doxycycline (DOX), Sulphonamides cp. (SUL), Trimethoprim (TMP), Azithromycin (AZM), Streptomycin (STR), Tobramycin (TOB), Amikacin (AMK)

**Table S2** Detailed list of genetic changes in chromosomes of plasmid-free strains in comparison to their wild-type isolates.

| Isolate ID | Plasmid-free clone ID[a] | Number of SNPs[b] | Non-synonymous SNPs[b] | Synonymous SNPs[b] |
|---|---|---|---|---|
| Dog168 | **CC4, CC5** | 0 | - | - |
| | CC1 | 1 | Ribonuclease E | - |
| | CC6 | 4 | Ribosomal protein S6 - L-glutamine ligase<br>YeiG formylglutathione hydrolase<br>RNA polymerase subunit beta R687C<br>RNA polymerase subunit beta D516G | - |
| OV24 | **CC2** | 0 | - | - |
| | CC1, CC4 | 1 | DNA polymerase III subunit tau | - |
| M45 | **CC2, CC3, CC4** | 0 | - | - |
| | CC1 | 1<br>1 | YohK (inner membrane protein)<br>ZntB zinc transport | - |
| M24 | **CC5, CC8** | 0 | - | - |
| | CC6 | 2 | Selenocysteine-specific elongation factor | PTS system EIIBCA component |
| | CC7 | 2 | Ribonuclease E | Fimbria adhesin EcpD |
| M70 | **CC1** | 0 | - | - |
| | CC2 | 3 | Valine--tRNA ligase<br>Lipopolysaccharide core heptose kinase RfaP<br>Outer membrane usher protein FimD | - |
| | CC3 | 3 | SrpC efflux pump | intergenic<br>arylsulfatase |
| | CC4 | 6 | NanR transcriptional repressor<br>DNA polymerase I<br>L-threonine dehydrogenase<br>Autotransporter outer membrane beta-barrel domain-containing protein | intergenic<br>fhuF ferric iron reductase protein |

Plasmid-free clones highlighted in bold were selected for fitness experiments

[a]IDs of the plasmid-free clones, CC stands for cured clone; [b]SNPs stands for single nucleotide mutations

#### References


1. Jamborova I, Johnston BD, Papousek I, et al. Extensive Genetic Commonality among Wildlife, Wastewater, Community, and Nosocomial Isolates of *Escherichia coli* Sequence Type 131 (*H30*R1 and *H30*Rx Subclones) That Carry *bla*$_{CTX-M-27}$ or *bla*$_{CTX-M-15}$. *Antimicrob Agents Chemother*. 2018;62(10):1-15.
